# Low-dose inhalation exposure to trichloroethylene induces dopaminergic neurodegeneration in rodents

**DOI:** 10.1101/2023.07.12.548754

**Authors:** Ashley B. Adamson, Neda M. Ilieva, William J. Stone, Briana R. De Miranda

**Affiliations:** Center for Neurodegeneration and Experimental Therapeutics, Department of Neurology, University of Alabama at Birmingham, Birmingham, AL, USA

## Abstract

Trichloroethylene (TCE) is one of the most pervasive environmental contaminants in the world and is associated with Parkinson disease (PD) risk. Experimental models in rodents show that TCE is selectively toxic to dopaminergic neurons at high doses of ingestion, however, TCE is a highly volatile toxicant, and the primary pathway of human exposure is inhalation. As TCE is a highly lipophilic, volatile organic contaminant (VOC), inhalation exposure results in rapid diffusion throughout the brain, avoiding first-pass hepatic metabolism that necessitated high doses to recapitulate exposure conditions observed in human populations. We hypothesized that inhalation of TCE would induce significantly more potent neurodegeneration than ingestion and better recapitulate environmental conditions of vapor intrusion or off gassing from liquid TCE. To this end, we developed a novel, whole-body passive exposure inhalation chamber in which we exposed 10-month-old male and female Lewis rats to 50 ppm TCE (time weighted average, TWA) or filtered room air (control) over 8 weeks. In addition, we exposed 12-month-old male and female C57Bl/6 mice to 100 ppm TCE (TWA) or control over 12 weeks. Both rats and mice exposed to chronic TCE inhalation showed significant degeneration of nigrostriatal dopaminergic neurons as well as motor and gait impairments. TCE exposure also induced accumulation of pSer129-αSyn in dopaminergic neurons as well as microglial activation within the substantia nigra of rats. Collectively, these data indicate that TCE inhalation causes highly potent dopaminergic neurodegeneration and recapitulates some of the observed neuropathology associated with PD, providing a future platform for insight into the mechanisms and environmental conditions that influence PD risk from TCE exposure.

## Introduction

Trichloroethylene (TCE) is a chlorinated organic solvent used as a degreasing product and as chemical feedstock for developing refrigerants ^1,2^. Due to its widespread use over much of the 20^th^ century, TCE is now a pervasive environmental contaminant in the United States and throughout the world ^3^. TCE has been measured in 9-34% of drinking water within the United States ^4^, and non-aqueous plumes of TCE within groundwater aquifers are resistant to degradation ^5^, presenting a challenge for both tracking and mitigating TCE exposures ^6,7^. Due to its highly volatile nature, the most common route of exposure in humans is through inhalation ^8,9^, which can readily occur from contaminated air or volatilization from groundwater and soil ^10,11^. As the vaporization point for TCE is 70°F (21°C), vapor intrusion into indoor air of businesses and homes represents an environmental risk for TCE exposure ^8,10^, which can be impacted by indoor and outdoor temperature, humidity, and a number of other dynamic physical properties ^12-14^. Consumer products are also primary sources for indoor air contamination of chlorinated volatile organic chemicals (VOCs), and TCE is within the top 15 highest estimated emissions of Prop 65-listed chemicals in consumer products ^15^. As TCE is a known carcinogen, several agencies regulate limits of exposure in air and water (summarized in **Table 1**).

**Table 1.**
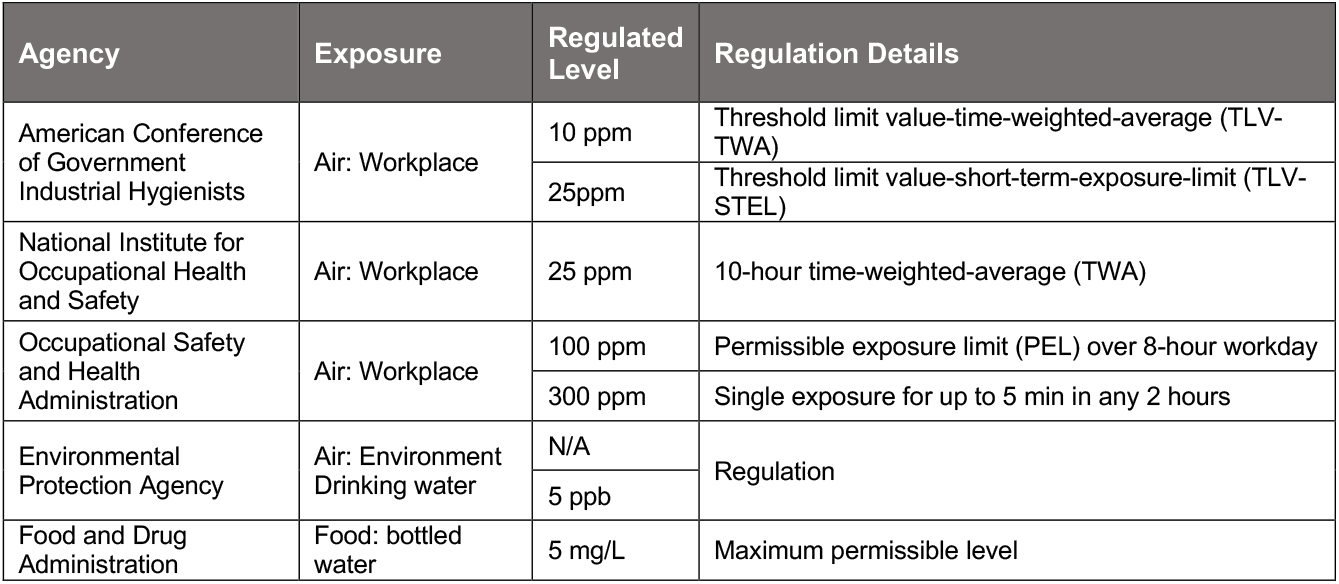
Summary of TCE exposure regulations from US institutions.

| Agency | Exposure | Regulated Level | Regulation Details |
| --- | --- | --- | --- |
| American Conference of Government Industrial Hygienists | Air: Workplace | 10 ppm | Threshold limit value-time-weighted-average (TLV-TWA) |
|  |  | 25ppm | Threshold limit value-short-term-exposure-limit (TLV-STEL) |
| National Institute for Occupational Health and Safety | Air: Workplace | 25 ppm | 10-hour time-weighted-average (TWA) |
| Occupational Safety and Health Administration | Air: Workplace | 100 ppm | Permissible exposure limit (PEL) over 8-hour workday |
|  |  | 300 ppm | Single exposure for up to 5 min in any 2 hours |
| Environmental Protection Agency | Air: Environment<br>Drinking water | N/A | Regulation |
|  |  | 5 ppb |  |
| Food and Drug Administration | Food: bottled water | 5 mg/L | Maximum permissible level |

TCE is linked to elevated risk for Parkinson’s Disease (PD) ^18-22^, with growing evidence including a large case-control study of veterans in Camp Lejeune, North Carolina (odds ratio, 1.70; 95% CI, 1.39-2.07; P < .001) ^23^. We and others have shown that systemic exposure to TCE in rodents induces the selective neurodegeneration of dopamine neurons in the substantia nigra (SN) and their terminal projections in the striatum (ST), accumulation of endogenous α-synuclein (αSyn), and microglial activation ^20,21,24-27^. However, previous studies investigating neurodegeneration caused by TCE have used oral gavage or intraperitoneal injection of high doses of TCE in rodents ^20,26,27^, which limits the translational relevance to most environmental exposure conditions. Inhalation exposure may particularly influence neurodegeneration as inhaled toxicants have a direct route into the brain through olfactory pathways ^28^, and from perfused blood from the lungs ^29^.

TCE is a small, highly lipophilic molecule that readily crosses membranes, including the blood-brain barrier, and causes acute neurological effects following inhalation such as dizziness, sleepiness, and reduced reaction time ^8,30,31^. We predicted that inhalation exposure would be especially toxic to dopaminergic neurons and produce Parkinsonian pathology at doses lower than previously used in other routes of exposure. To assess this, we developed an inhalation exposure platform using a whole-body, passive exposure inhalation chamber, coupled with environmentally relevant extrapolated doses, and assessed PD-related neurodegeneration through the most common route of TCE exposure in humans. We found that exposure to a time weighted average (TWA) of 50 ppm TCE for rats and 100 ppm TCE for mice, over 8 and 12 weeks respectively, resulted in significant loss of nigrostriatal dopaminergic neurons that corresponded with motor behavior deficits. We also observed accumulation of phosphorylated αSyn (pSer129-αSyn) within dopaminergic neurons as well as activated microglia within the ventral midbrain. Collectively, we found that TCE inhalation resulted in pathologic outcomes consistent with idiopathic PD pathology in human brain tissue, providing further support for epidemiological data that TCE is a risk factor for PD.

## Materials and Methods

### Chemical reagents and supplies

TCE (CAS 79-01-6) and other chemicals were purchased from Sigma-Aldrich (St. Louis, MO) unless otherwise noted. Antibody information for immunohistochemistry (IHC) is listed in Table 2.

**Table 2.** Antibody information for brain tissue immunohistochemistry.

| Antigen | Antibody Catalog # | Company | Immunohistochemistry Concentration |
| --- | --- | --- | --- |
| Tyrosine hydroxylase | AB512 | EMD Millipore (Burlington, MA) | 1:2000 |
| Ser129-α-Synuclein | ab51253 | Abcam (Cambridge, MA) | 1:500 |
| IBA1 | 019-19741 | Wako Chemical USA (Irvine, CA) | 1:500 |
| CD-68 | MCA341 | BioRad (Hercules, CA) | 1:500 |
| NeuroTrace | N21483 | Invitrogen (Waltham, MA) | 1:500 |

### Animals

Adult (10-month-old) male and female Lewis rats were obtained through a retired breeding program from Envigo (Indianapolis, IN) and adult (12-month-old) male and female C57BL/6 mice were obtained from Jackson Laboratory (Bar Harbor, ME). Upon arrival, rats and mice were acclimated to the UAB small animal facility for one week prior to study onset. Rodents were provided with standard rodent chow and filtered water ad libitum throughout the study period. Rats were double housed with platforms and wooden blocks for enrichment, mice were housed 3-4 to a cage with enrichment nests, and all rodents maintained at 72-74°C with 50-60% humidity. At the study termination, rats and mice were humanely euthanized using a lethal dose of pentobarbital euthanasia and underwent transcardial perfusion with PBS, followed by organ collection. All animal experiments were conducted with approval by the University of Alabama at Birmingham Institutional Animal Care and Use Committee (IACUC).

### TCE administration

TCE and control (HEPA filtered room air) groups were randomly divided. TCE exposed rodents were placed in a whole-body passive exposure chamber inside a chemical fume hood within the lab. Control animals were placed in an adjacent fume hood with passive flow of filtered room air. Rats were exposed for 7 hours a day for 8 weeks, and mice were exposed 7 hours a day for 12 weeks (to compensate for metabolism rate differences). TCE concentrations were monitored using Honeywell MiniRAE3000 photoionization detector (PID monitor) placed at the level of the rat/mouse, equipped with a 11.7 UV lamp using a correction factor of 0.43 for trichloroethylene. TCE handling and disposal was carried out following UAB Environmental Health and Safety procedures for hazardous chemicals.

### Motor Behavior

Motor behavior was assessed using Noldus Catwalk XT 10.6 gait analysis system in the UAB Behavioral Assessment Core. Rats and mice were trained to walk across the Catwalk stage one day prior to testing. The camera distance was set at 70 cm for rats and 35 cm for mice. Each subject required three successful runs for analysis. A successful run was defined as a duration from 0.5-5.0 seconds with a maximum variation of 60%. Detection settings were automatically generated, and gain was varied from 220-225 to optimize signal produced from pawprints. During testing, animals were allowed to walk across the stage, and pawprints were recorded. Animals were excluded after 30 failed run attempts. Prints were analyzed in Catwalk XT software (version). Compliant runs were automatically classified, manually validated, and extraneous signal prints were excluded manually while blinded.

### Striatal terminal intensity

Serial brain sections (35 μm) spanning the volume of the rat and mouse striatum (one-sixth sampling fraction, approximately 10 sections per animal) were stained for tyrosine hydroxylase (TH) and detected using an infrared secondary antibody (IRDye 680, Licor Biosciences). Striatal tissue sections were analyzed using near-infrared imaging for density of dopamine neuron terminals (LiCor Odyssey) and analyzed using LiCor Odyssey software (V3.0; Licor Biosciences, Lincoln, NE). Results are reported as striatal TH intensity in arbitrary fluorescence units.

### Stereology

Stereological analysis of dopamine neuron number in the substantia nigra (SN) was adapted from Cao et al 2010 employing an unbiased, automated system^32^. Briefly, serial nigral tissue sections (one-sixth sampling fraction, 10 sections per animal) spanning the volume of the SN were stained for TH and quantified using a MBF Biosciences Widefield microscope and Stereo Investigator software (Center for Neurodegeneration and Experimental Therapeutics Core, UAB). Representative fluorescent images were processed using Nikon NIS-Elements Advanced Research software (Version 4.5, Nikon, Melville, NY), and quantitative analysis was performed on brightfield images. Results are reported as the number of TH-positive cell bodies (whole neurons) within the SN.

### Immunohistochemistry

Brain sections were maintained at −20°C in cryoprotectant, stained while free-floating, and mounted to glass slides for imaging. Fluorescent immunohistochemical images were collected using a Nikon AX confocal microscope. Quantitative fluorescence measurements and all imaging parameters were kept consistent across specimens. Confocal images were analyzed using Nikon NIS-Elements Advanced Research software (Version 4.5, Nikon, Melville, NY). A minimum of 4 images per tissue slice were analyzed per animal, averaging 9–15 neurons per 60x image. Results are reported as a measure of puncta within TH-positive cells, either number of objects (# of objects) or area (μm^2^).

### Statistical analysis

All data were expressed as mean values ± standard error of the mean. Outcome measures were evaluated for normally distributed means using a normality test for homoscedasticity, followed by appropriate parametric or non-parametric tests; student’s unpaired two-tailed t-test to compare control and TCE. Statistical significance between male and female rats was evaluated for normally distributed means by two-way analysis of variance (two-way ANOVA) with a Sidak multiple comparisons test to correct for mean comparison between multiple groups, where source of variation was defined as “sex” or “TCE” groups. Prior to study onset, an a priori power analysis was conducted using G*power software to determine the sample size required for a 20–40% difference between mean, with a 95% power at α = .05. Statistical significance between treatment groups is represented in each figure as *p < .05, **p < .01, ***p < .001, ****p < .0001, unless otherwise specified on graph. Statistical outliers from each data set were determined using the extreme studentized deviate (Grubbs’ test, α = .05). Statistical analyses were carried out using GraphPad Prism software (V. 5.01). Animal treatment groups obscured and renumbered by a third party. Animals were unblinded at the conclusion of the analysis.

## Results

### Establishing human equivalent dose (HED)

To model a low HED of TCE in both rats and mice, we used allometric scaling to normalize TCE dose to body surface area to ascertain the animal equivalent dose (AED) of TCE. Our HED was set at 8 ppm, 92 ppm lower than currently regulated OSHA permissible exposure (Table 1) for TWA. Rat and mouse correction factors (6.2 and 12.3 respectively) were obtained from Nair et al ^33^ to scale body surface area for a 60 kg human. As allometric scaling does not encompass differences in the metabolism of TCE, the exposure window was increased for mice to accommodate their increased metabolic rate ^30^. Our AED for rats and mice was obtained through the following equation:

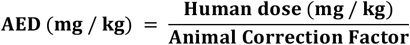

TCE concentration was a TWA of 50 ppm for rats and 100 ppm for mice and fluctuations were continuously recorded on a PID monitor (Fig. 1).

**Figure 1.**
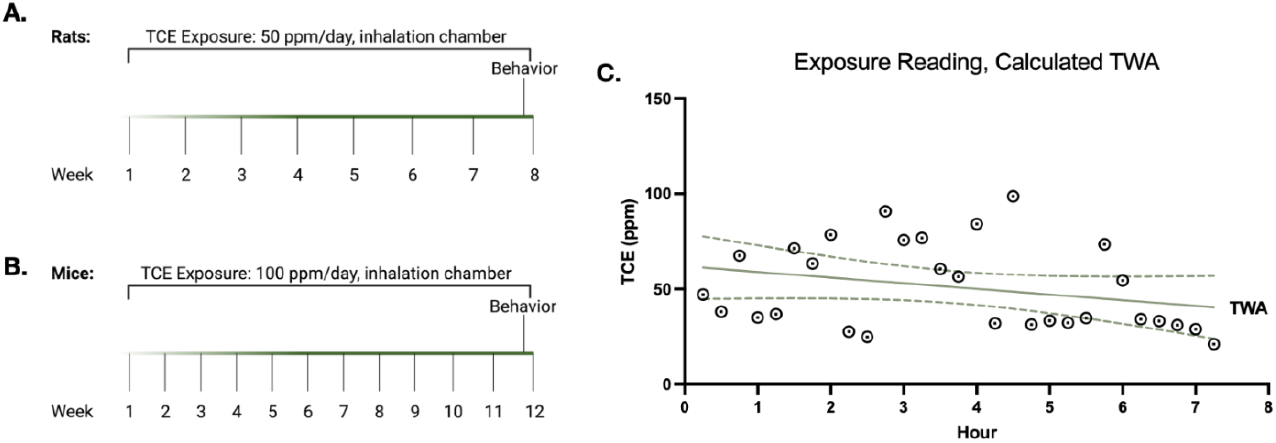
Chronic TCE inhalation exposure in adult rats and mice. (**A**) 10-month-old male and female Lewis rats were exposed to 50 ppm time-weighted average (TWA) of TCE inhalation for 7 hours per day, 5 days per week, for 8 weeks. (**B**) 12-month-old male and female C57Bl/6 mice were exposed to 100 ppm TWA of TCE inhalation for 7 hours per day, 5 days per week, for 8 weeks. (**C**) Representative plot showing TCE concentration changes in 15-minute increments over 7 hours for a 50 ppm TWA.

### TCE inhalation exposure induced nigrostriatal dopaminergic neurodegeneration and motor deficits in rats

Immunohistochemistry (IHC) of striatal brain sections from TCE-exposed 10-month-old make and female Lewis rats revealed over a 50% reduction in tyrosine hydroxylase (TH) intensity compared to control (p < 0.0001; Fig. 2 A, C). In addition, TCE exposure caused a loss of approximately 50% of dopaminergic neurons assessed by unbiased stereology in the SN compared to control (p = 0.0003; Fig. 2 B, D). The loss of neurons was confirmed by staining all cells with NeuroTrace, a fluorescent Nissl dye, which showed significant cell loss following TCE exposure (Supplemental Fig. 1). Following 8 weeks of 50 ppm TCE inhalation exposure, male rats were assayed for motor behavior deficits utilizing Noldus Catwalk XT gait analysis system, which revealed a number of affected gait parameters. Left swing speed (p = 0.0012), front swing speed (p = 0.0350), and hind limb swing speed (p = 0.0040) and hind print area (p = 0.0041) were increased following TCE exposure when compared to vehicles. Conversely, front stand time (p = 0.0437), left (p = 0.0479) and front step cycle (p = 0.0400), and hind base of support (p = 0.0453) were decreased in TCE exposed animals compared to controls (Fig. 2 E-N). In all collated limb analyses, the deficits appear to be driven by left side limbs indicating an asymmetric gait impairment caused by exposure to TCE inhalation.

**Figure 2.**
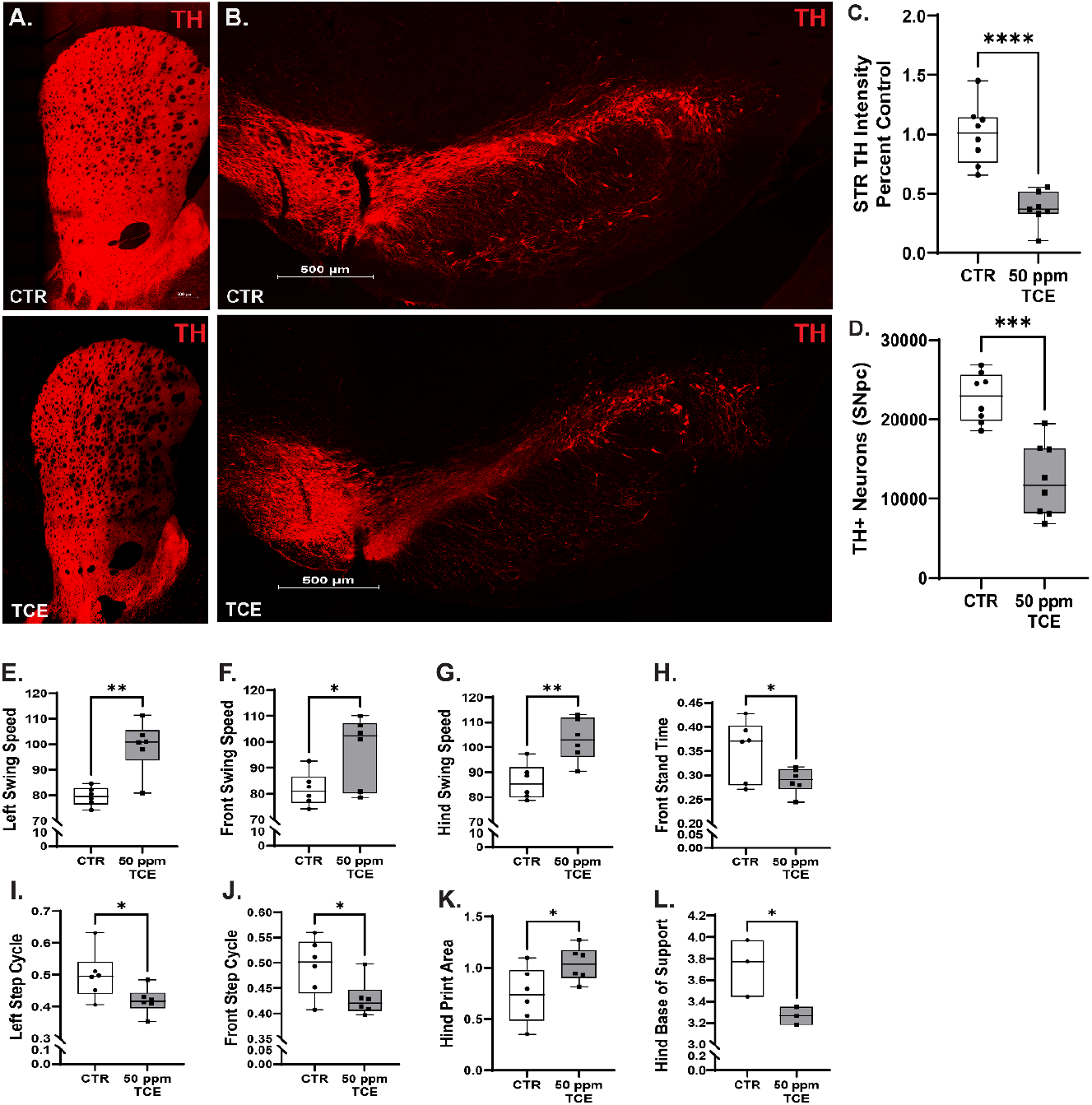
Inhalation exposure to TCE induces nigrostriatal dopaminergic neurodegeneration and motor deficits in adult Lewis rats. Representative images (20x) of 35 μm brain tissue sections of the striatum (**A**) and substantia nigra (**B**) immunostained for tyrosine hydroxylase (TH) from male and female Lewis rats exposed to 50 ppm TCE inhalation or filtered room air (control). Quantification of dopaminergic terminal loss from the striatum (**C**) and dopaminergic neuron loss from the SNpc (**D**). Quantitative parameters measured from the Noldus CatWalk XT gait analysis system showed significant differences in left swing speed (p = 0.0012; **E**), front swing speed (p = 0.0350; **F**), hind swing speed (p = 0.0040; **G**), front stand time (p = 0.0437; **H**), left step cycle (p = 0.0479; **I**), front step cycle (p = 0.0400; **J**), hind print area (p = 0.0041; **K**), and hind base of support (p = 0.0453; **L**). Statistical analysis unpaired t-test, error bars represent SEM, (N = 8).

### TCE inhalation increases pSer129-αSyn accumulation in dopaminergic neurons and induces microglial activation within the substantia nigra of exposed rats

Endogenous pSer129-αSyn was assayed within dopaminergic (TH+) neurons using IHC and quantified by accumulated puncta. TCE inhalation caused significant accumulation of endogenous pSer129-αSyn compared to control (p = 0.0049; Fig. 3), which was predominantly located within the soma and proximal axons of dopaminergic neurons of rats exposed to TCE. In addition, rats exposed to 50 ppm TCE via inhalation displayed increased CD68 expression, a lysosomal protein and macrophage phagocytic activity marker, within IBA1 positive cells compared to control (p = 0.0117; Fig. 4). Microglia also displayed an activated morphological phenotype within TCE exposed rat brain tissue, appearing more hypertrophic with less ramified processes and amoeboid shape.

**Figure 3.**
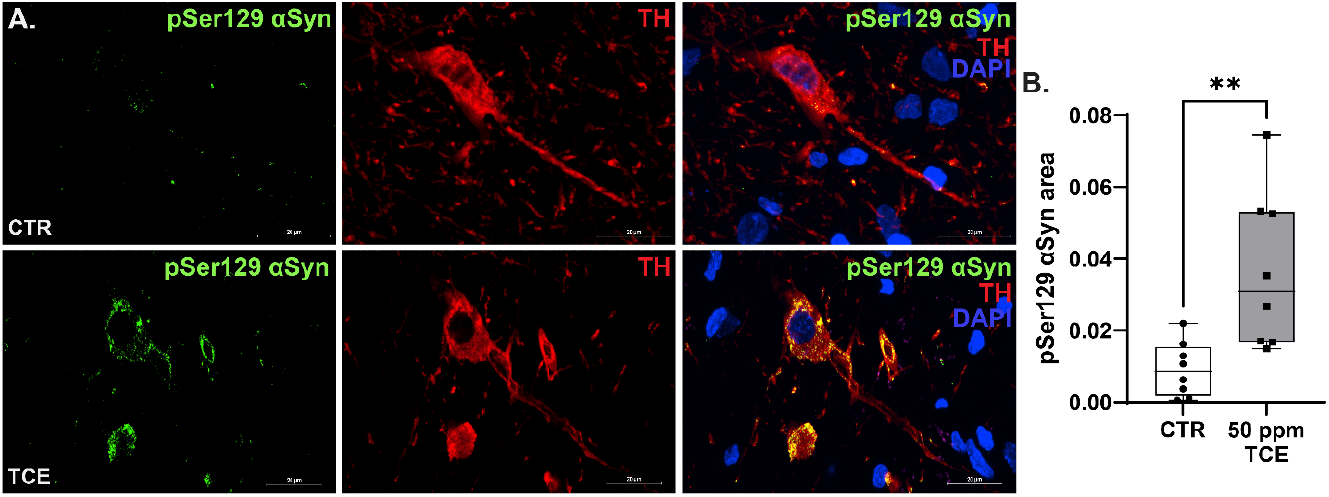
TCE inhalation exposure induced the accumulation of pSer129-αSyn in dopaminergic neurons. (**A**) Representative confocal images (60x) of pSer129-αSyn expression (green) within dopaminergic neurons immunoreactive for tyrosine hydroxylase (red). (**B**) Quantification of pSer129-αSyn area within dopaminergic neurons following TCE exposure (p = 0.0049). Statistical analysis unpaired t-test, error bars represent SEM, (N = 8 vehicle, 8 TCE).

**Figure 4.**
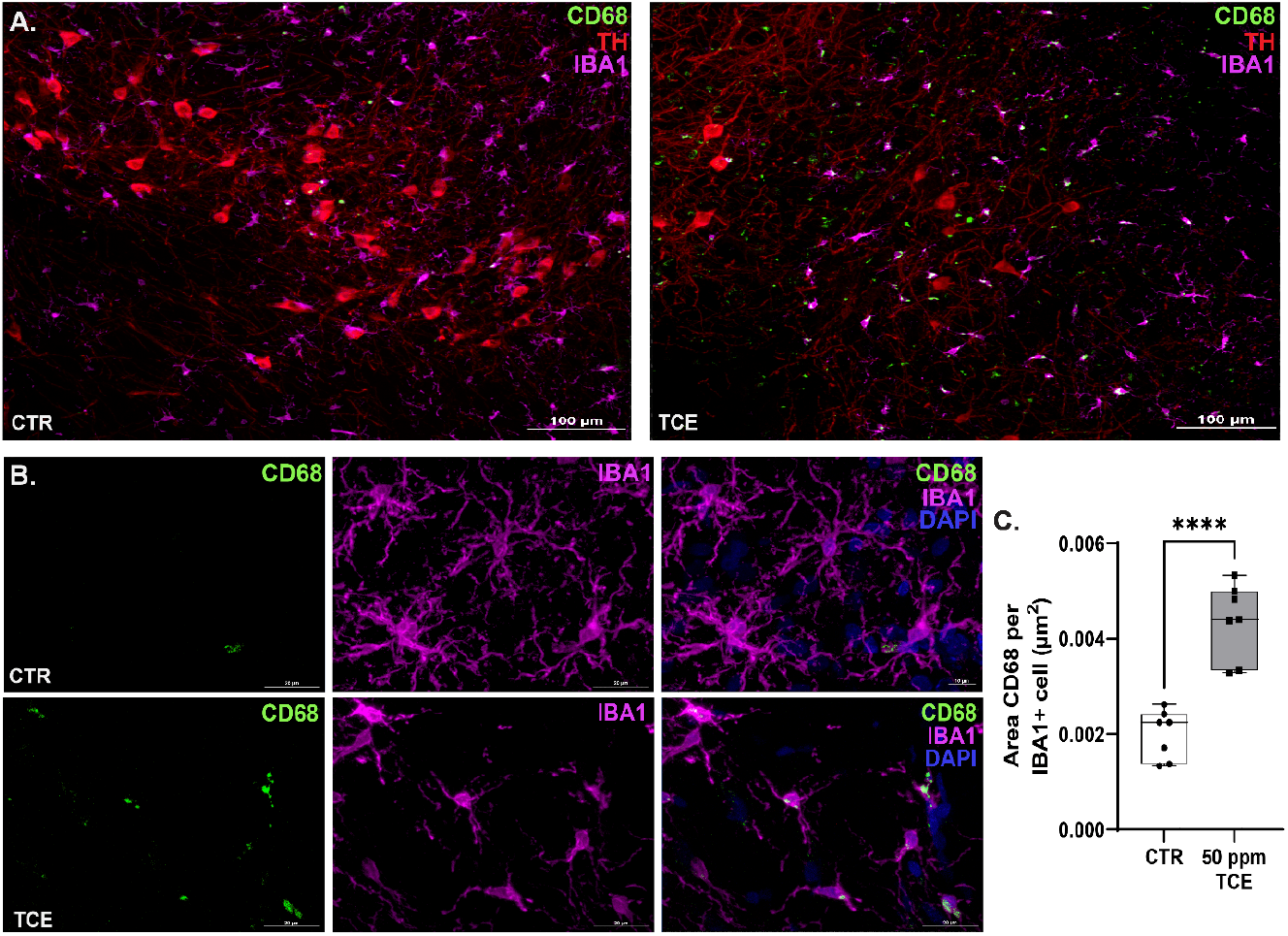
Microglial activation was observed in the substantia nigra of adult rats exposed to TCE via inhalation. (**A**-**B**) Representative confocal images (20x and 60x) of phagocytic activation marker CD68 (green) within microglia immunoreactive for IBA1 in the SN (magenta). (**C**) Quantification of CD68 area within microglia following TCE exposure (p = 0.0117). Statistical analysis unpaired t-test, error bars represent SEM, (N = 8 vehicle, 8 TCE).

### Sex differences from TCE inhalation were not observed in outcome measures

Age-matched adult male and female Lewis rats (10 months old) were obtained from the same commercial source (Envigo) and housed under identical conditions prior to the onset of TCE exposure; 50 ppm of TCE over 8 weeks. Histopathological analysis of the nigrostriatal tract showed no significant differences between sexes with loss of dopaminergic neuron terminals or somas of TCE exposed rats (p = 0.8959; p = 0.0868; Fig. 5 A-B). Additionally, no differences were found between sexes in TCE exposed groups in the expression of pSer129-αSyn in the SN (p = 0.9180; Fig. 5 C). Finally, in TCE animals, no sex differences were apparent in the expression of CD68 within IBA1 positive cells (p = 0.5115; Fig. 5 D).

**Figure 5.**
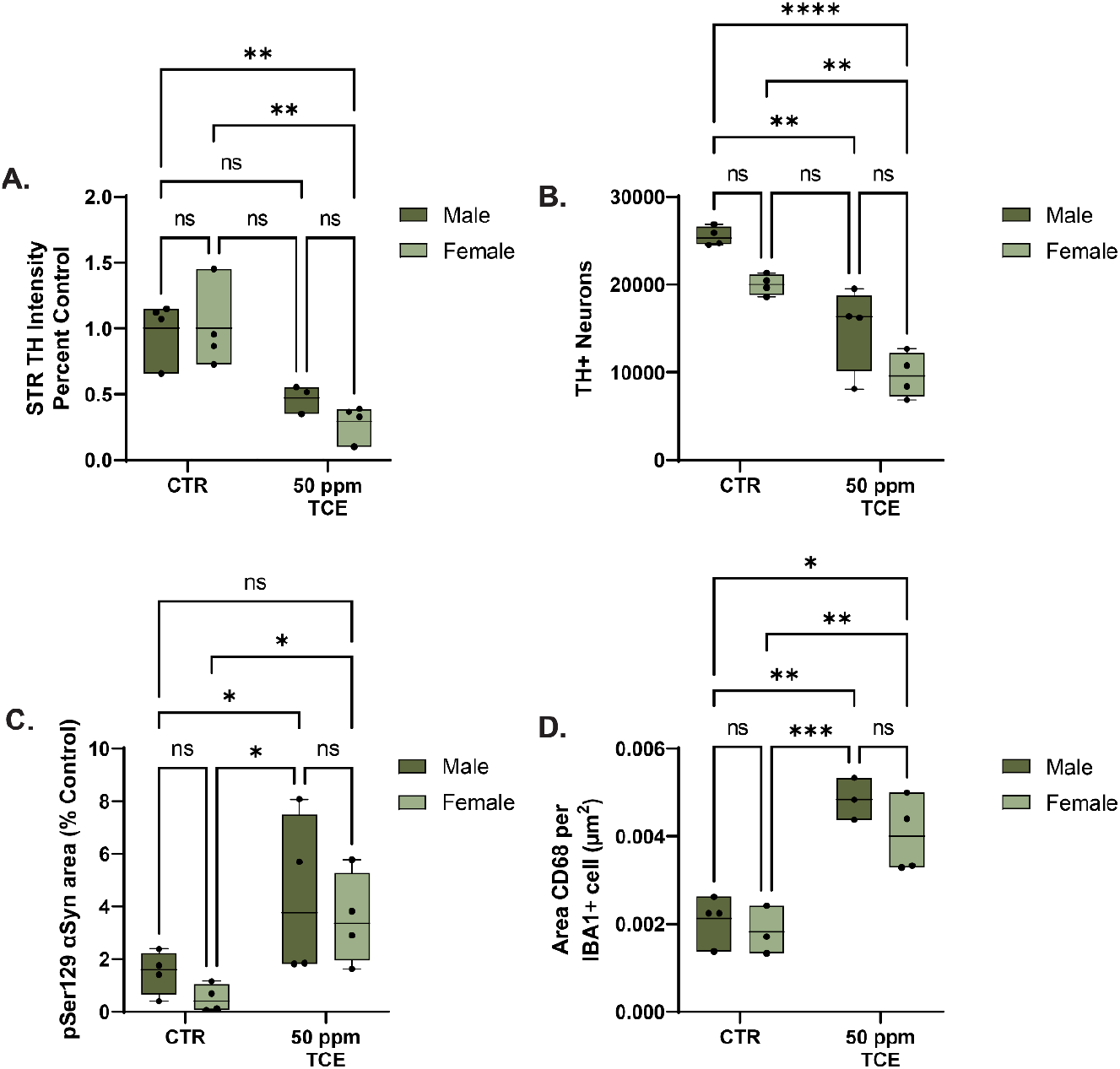
Sex differences in cellular pathology were not observed in male and female rats exposed to inhaled TCE. (**A**) Quantitative analysis of striatal TH intensity represented as percent of control subjects indicated no significant differences caused by sex of the animals (p = 0.8959). (**B**) Quantification of dopaminergic neurons did not significantly differ between sexes (p = 0.0868). (**C**) pSer129-αSyn expression of TCE exposed animals did not differ between sexes (p = 0.9180). (**D**) Quantitative analysis of activated microglia showed no difference in CD68 staining between sexes (p = 0.5115). Statistical analysis two-way ANOVA with Fisher’s Least Significant Difference, error bars represent SEM, (N = 4 animals/sex/group).

### TCE inhalation induced nigrostriatal dopaminergic neurodegeneration and motor deficits in mice

Adult (12 month) C57Bl/6 male and female mice were exposed to 100 ppm TCE inhalation of vehicle (HEPA filtered room air). Immunohistochemistry of striatal brain sections from TCE-exposed mice had approximately a 30% reduction in TH intensity when compared to vehicles (p < 0.0071; Fig. 6 A, C). In addition, TCE exposure caused a loss of approximately 50% of dopaminergic neurons in the SN when compared to vehicles (p = 0.0391; Fig. 6 B, D). The loss of neurons was confirmed by staining all cells with NeuroTrace which showed significant cell loss following TCE exposure (Supplemental Fig. 1). Following 12 weeks of TCE exposure, motor behavior was assayed using Noldus CatwalkXT gait analysis system. Mice exposed to TCE had a significantly increased left side swing speed (p = 0.0220) and reduced right side step cycle when compared to vehicles (p = 0.0221; Fig. 6 E-H). As observed in rats, these findings indicate an asymmetric gait impairment commonly seen in individuals with PD^34^. These results indicate that this exposure method can be applied in both rat and mouse models of disease.

**Figure 6.**
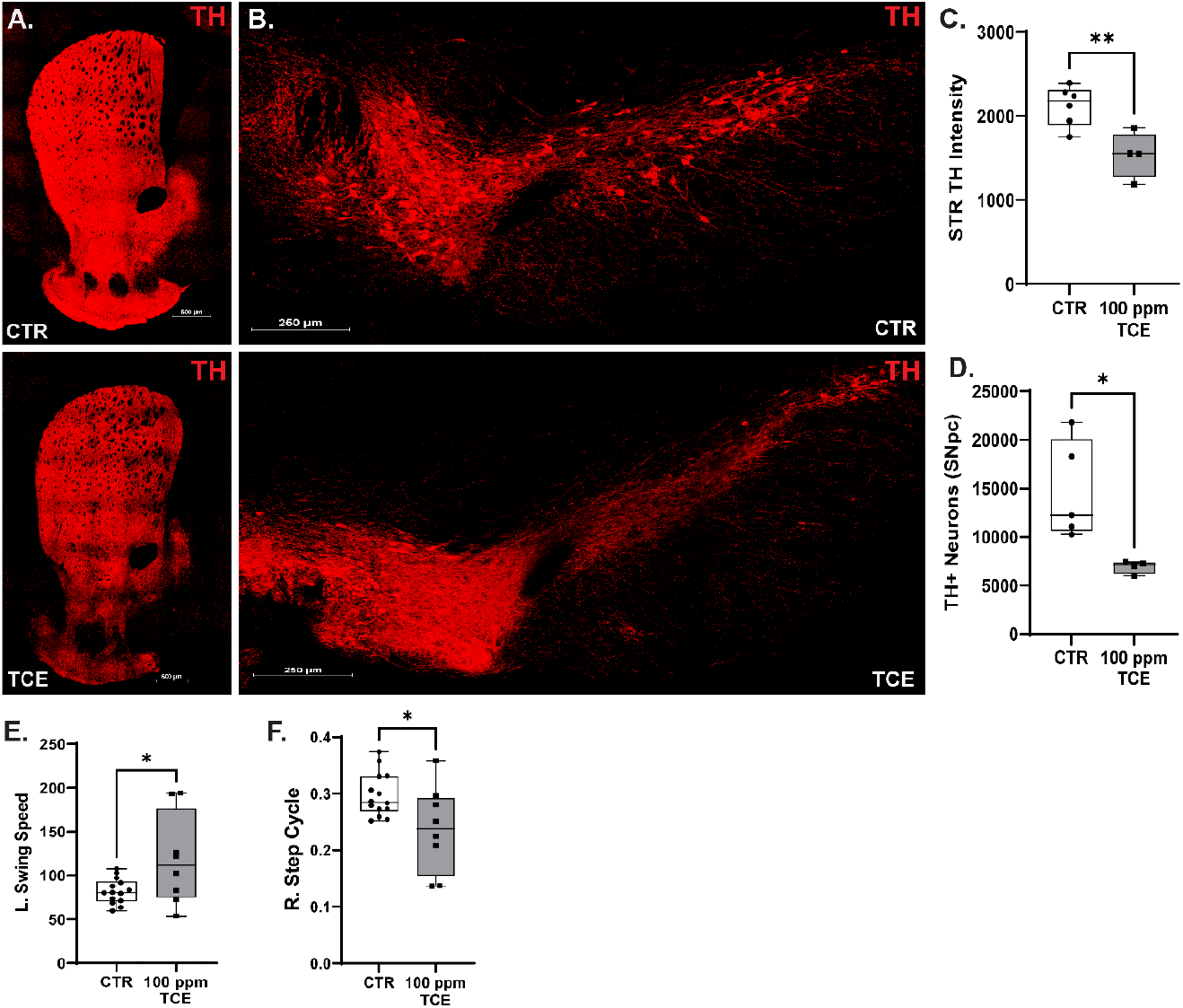
TCE inhalation exposure caused nigrostriatal dopaminergic neurodegeneration and gait abnormalities in wildtype mice. Representative montage images (20x) of 35 μm brain tissue sections of the striatum (**A**) and substantia nigra **(B**) immunostained for tyrosine hydroxylase (TH) from male and female C57Bl/6 mice exposed to 100 ppm TCE inhalation or filtered room air (control) for 12 weeks. Quantification of dopaminergic terminal loss from the striatum (p = 0.0071) (**C**) and dopaminergic neuron loss from the SNpc (p = 0.0194) (**D**) showed significant TH loss (N=4-5). Quantitative parameters measured from the Noldus CatWalk XT gait analysis system showed significant differences in left swing speed (p = 0.0220) (**E**) and right step cycle (p = 0.0221) (**F**) of TCE exposed mice. Statistical analysis unpaired t-test, error bars represent SEM, (N = 14 vehicle, 8 TCE).

## Discussion

Growing evidence suggests that TCE exposure elevates risk for PD ^18,35^, however, considerably less is known about how this pervasive environmental contaminant causes the specific neurotoxicity that drives parkinsonian neurodegeneration. The most common route of human exposure to TCE is from inhalation of its vapor, which typically occurs through occupational use of the solvent or from vapor intrusion into building airspace from contaminated soil and/or groundwater ^8-11^. Previously reported experimental data shows that TCE exposure in rodents induces the selective degeneration of dopaminergic neurons from the nigrostriatal tract, as well as other PD-related pathologies such as motor deficits, neuroinflammation, and pSer129-αSyn accumulation ^20,21,24-27^. However, the majority of this evidence was generated from exposure through oral gavage administration, which represents a less common TCE exposure route (ingestion) and requires higher doses of TCE in rats and mice than what is typically recorded in human populations.

For example, in both Gash et al. (2008) and Liu et al., (2010) 1,000 mg/kg TCE was administered via oral gavage over 6 weeks in Fisher 344 rats to induce significant dopaminergic neurodegeneration ^21,26^. In C57Bl/6 mice, 400 mg/kg TCE over 8 months induced a similar severity of nigral dopaminergic cell loss as reported in rats, and also caused motor impairments ^37^. With interest in examining moderate experimental doses, our group reported in 2021 that daily oral gavage with 200 mg/kg TCE over 6 weeks in aged Lewis rats (10-15 months old) caused significant loss of nigrostriatal dopaminergic neurons, induced endolysosomal dysfunction and αSyn aggregation, caused neuroinflammation, and influenced aberrant kinase activation of the PD-associated protein LRRK2^38^.

As a comparison for TCE dose in animals, the most well-known drinking water contamination event recorded in the US stems from Camp Lejeune, North Carolina, where reconstruction data estimates an average of 783 μg/L TCE, with an actual measured maximum value of 1,400 μg/L ^39^. These levels of exposure to a 60 kg human would be equivalent to approximately 0.62-1.23 mg/kg daily TCE in a rat or mouse, respectively ^33^. However, exposure to contaminated water at Camp Lejeune occurred for over 30 years, varied by time, season, and location, and included other toxicants such as tetrachloroethylene and benzene ^39^, all of which are conditions that cannot be fully recapitulated in laboratory animals. Likewise, exposure to TCE at Camp Lejeune, as in other contaminated sites, included inhalation of TCE from vaporization of contaminated groundwater and soil, ingestion, and dermal exposure from TCE-contaminated drinking water. In addition, interspecies variability in the oxidative metabolism of TCE indicates that rodents metabolize TCE more rapidly than humans ^40^. Mice even exhibit strain-dependent variability of metabolism in ^41-44^, suggesting that greater doses beyond simple allometric scaling would be necessary in laboratory animals to accurately replicate exposure. Thus, dose conversions of TCE exposure from animal to human and vice versa, are complex, rely on imperfect conversion assumptions, and should be considered in the temporal context of a human lifespan. Toward this end, inhalation provides an environmentally relevant route of exposure with the potential for chronic timepoints that more closely match conditions observed in TCE-exposed populations. Here, we selected an inhalation exposure of 50 ppm TCE in rats and 100 ppm TCE in mice (TWA) – doses that are at or below regulated occupational exposure limits in humans (**Table 1**), and though non-exact, allometrically scale to a HED of ∼8 ppm ^33^.

While this was not the first study to examine TCE inhalation as a primary route of exposure in rodents, to our knowledge, this is the first data on parkinsonian neurodegeneration or associated pathology from passive, chronic TCE inhalation. Previous studies investigating the effects of TCE inhalation on renal toxicity, carcinogenicity, neuronal plasticity in hippocampal and visual cortical slices, and the organization of motor behavior have been reported; however, doses used in these studies were often high, in some cases up to 1500 ppm, and were predominantly acute exposures ^45-48^. Given the selective sensitivity of dopaminergic neurons to neurotoxicants ^49^, we hypothesized that TCE inhalation would induce more severe neurodegeneration than ingestion, forgoing first-pass hepatic metabolism within the liver ^50^ and resulting in neurotoxicity at lower doses than previously used. In addition, exposure to inhaled toxicants presents more direct routes to the brain through the lungs as well as the olfactory bulb, from which some efferent pathways synapse directly in the striatum ^28,51^. Indeed, despite a lower dose, 50 ppm TCE inhalation caused more dopaminergic neurodegeneration than 200 mg/kg TCE via oral ingestion: ∼50% reduction in the number of dopaminergic neurons from TCE inhalation versus ∼35% from oral gavage ^24^. This may have partially been impacted by duration of exposure (8 weeks of 50 ppm TCE exposure versus 6 weeks of 200 mg/kg exposure); however, given the vast difference in TCE concentration between the two exposure methods, route of exposure rather than duration likely provided the most significant impact on neurodegeneration caused by TCE.

Route of exposure to environmental PD risk factors may also be a key variable in disease etiology. Prodromal symptoms of PD, such as anosmia and decreased gastrointestinal (GI) motility, can occur decades prior to the onset of motor dysfunction ^52-55^, and both olfactory and GI pathways are physiological entry points for environmental contaminants. Environmental influence on the Braak and dual-hit hypotheses, as predicated by Chen et al., (2022), suggests that PD may start in olfactory pathways, within the gut, or both, influenced by endogenous features like αSyn accumulation and exogenous factors such as toxicant exposures ^56^. TCE represents an environmental risk factor for PD that could influence disease pathogenesis through both inhalation (olfactory) and ingestion (gut) as it contaminates air and water. In line with this, we previously showed that ingestion of TCE caused gut microbiome changes that mirrored gut microbiome dysbiosis in idiopathic PD ^57^. Additionally, we observed the accumulation of pSer129-αSyn within dopaminergic neurons of TCE inhalation exposed rats, as well as microglial activation in the ventral midbrain, suggesting that αSyn dysfunction or inflammatory pathways induced by TCE could be potential triggers or facilitators of PD pathogenesis caused by exposure, which may be route dependent ^58^.

As previously discussed, species differences in TCE metabolism may also impact translational relevance to human exposure and disease. Published data in wildtype (C57Bl/6) mice show that both greater doses and longer exposure times were required to produce comparable dopaminergic neurodegeneration to rats ^37^. While rats may share more brain homology with humans than mice ^59^, mouse models encompass a broad scale of genetic manipulation that is not widely available in rat models, particularly for PD and other neurodegenerative disorders ^60^. To address this, we evaluated whether inhalation exposure to TCE in C57Bl/6 mice would produce similar dopaminergic pathology as observed in TCE exposed rats. To account for body surface area and metabolism differences, we adjusted the dose to 100 ppm TCE over 12 weeks, which induced comparable dopaminergic neurodegeneration (∼50% TH+ neuron loss) as well as asymmetric gait disturbances similar to those measured in rats. Thus, this model of TCE inhalation allows both the tissue homology of rats and genetic tools of mice to be leveraged in order to evaluate mechanisms of PD pathogenesis driven by TCE exposure, several of which are under current investigation in our lab.

As with all experimental models of exposure, there are limitations of TCE inhalation in rodents. For example, though inhalation may provide more direct brain access than ingestion, lung metabolism and elimination of TCE occurs rapidly ^50^, possibly resulting in different toxic metabolites based on species. In addition, though we did not observe sex differences at the level of outcome measures reported – mostly cellular pathological changes – this does not preclude sex-specific effects that could be apparent with more sensitive analyses, such as transcriptomic or epigenetic influences as found in other PD-related toxicant exposures ^61-63^. Finally, while nearly infinite combinations of exposure conditions could be investigated using this method, we chose to first report this proof-of-concept data in adult rodents with exposure occurring over 5 days per week, replicating a workplace or occupational exposure. Future studies to assess TCE exposure over the course of the lifespan, extremely low doses as observed in common environmental conditions, and the interaction with other PD risk factors will provide more information on the mechanisms of TCE-induced neurodegeneration.

## Conclusions

TCE has been implicated in PD risk for decades. However, although animal models of TCE exposure showed important proof of principle data of dopaminergic neurodegeneration and other PD related pathology, confusion over high doses required to recapitulate key features of the disease limited some translational relevance. We predicted this was predominantly because TCE exposure in humans mostly occurs via inhalation, providing a more direct path to the brain. Our data here suggest that TCE inhalation causes potent dopaminergic neurotoxicity at much lower doses than previously examined, providing a missing mechanistic link between TCE exposure and PD risk.

## Acknowledgments

This work was supported by research grants from the National Institutes of Environmental Health Sciences (R00ES029986, BRD), and the Parkinson Association of Alabama (BRD).

---

**Supplemental Figure 1.**
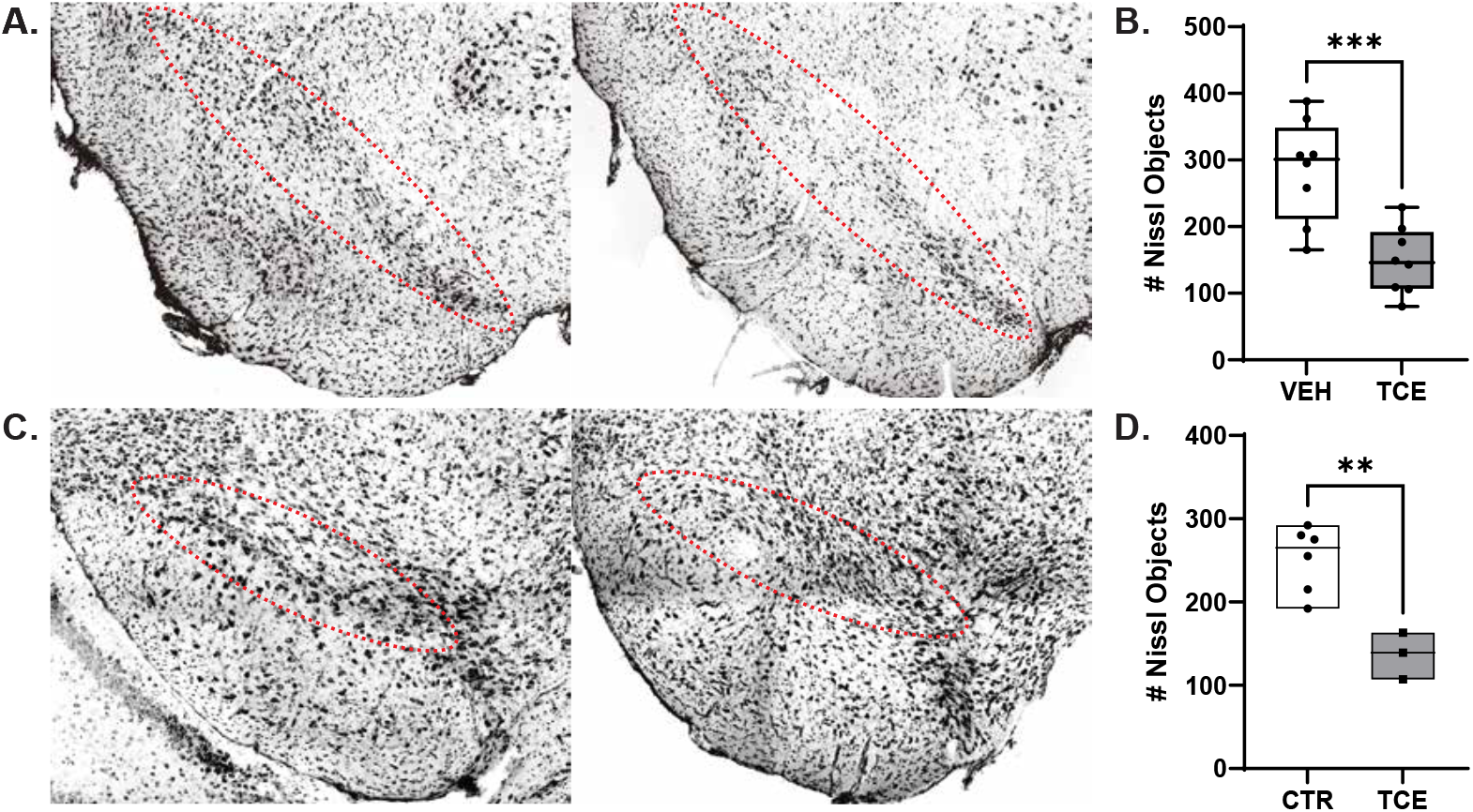
TCE exposure induces loss of Nissl+ cells in the substantia nigra of TCE exposed rats. Representative confocal images (20x) of Nissl-stained cells inside the boundaries of the SNpc shows loss of TH+ dopaminergic cell bodies in rats (**A**) and mice (**C**) exposed to 50 ppm or 100 ppm TCE respectively. Quantification of Nissl+ cells within the SN shows significant loss of cells in rats (p = 0.0009; **B**) and mice (p = 0.0030; **D**). Statistical analysis unpaired t-test, error bars represent SEM (N = 8 control, 8 TCE).

## Notes

### Competing Interest Statement

The authors have declared no competing interest.

## References

1. Doherty, R.E. (2000). A History of the Production and Use of Carbon Tetrachloride, Tetrachloroethylene, Trichloroethylene and 1,1,1-Trichloroethane in the United States: Part 1--Historical Background; Carbon Tetrachloride and Tetrachloroethylene. Environmental Forensics 1, 69–81. 10.1006/enfo.2000.0010.

2. IARC (1995). Trichloroethylene. IARC Monogr Eval Carcinog Risks Hum 63, 75–158.

3. Zogorski, J.S., Carter, J.M., Ivahnenko, T., Lapham, W.W., Moran, M.J., Rowe, B.L., Squillace, P.J., and Toccalino, P.L. (2006). Volatile organic compounds in the nation’s ground water and drinking-water supply wells. Report 1292. http://pubs.er.usgs.gov/publication/cir1292.

4. Epa (2000). Trichloroethylene. 2000/1//. https://www.epa.gov/sites/default/files/2016-09/documents/trichloroethylene.pdf.

5. Morrison, R.D., Murphy, B.L., and Doherty, R.E. (1964). 12 - Chlorinated Solvents. In Environmental Forensics, R.D. Morrison, and B.L. Murphy, eds. (Academic Press), pp. 259–277. https://doi.org/10.1016/B978-012507751-4/50034-3.

6. Wu, Y.J., Liu, P.G., Hsu, Y.S., Whang, L.M., Lin, T.F., Hung, W.N., and Cho, K.C. (2019). Application of molecular biological tools for monitoring efficiency of trichloroethylene remediation. Chemosphere 233, 697–704. 10.1016/j.chemosphere.2019.05.203.

7. Basu, N.B., Rao, P.S.C., Poyer, I.C., Annable, M.D., and Hatfield, K. (2006). Flux-based assessment at a manufacturing site contaminated with trichloroethylene. Journal of Contaminant Hydrology 86, 105–127. https://doi.org/10.1016/j.jconhyd.2006.02.011.

8. Todd, G.D., Ruiz, P., Mumtaz, M., Wohlers, D., Klotzbach, J.M., Diamond, G.L., Coley, C., and Citra, M.J. (2019). Toxicological profile for trichloroethylene (TCE).

9. Wu, C., and Schaum, J. (2000). Exposure assessment of trichloroethylene. Environ Health Perspect 108 Suppl 2, 359–363. 10.1289/ehp.00108s2359.

10. Archer, N.P., Bradford, C.M., Villanacci, J.F., Crain, N.E., Corsi, R.L., Chambers, D.M., Burk, T., and Blount, B.C. (2015). Relationship between vapor intrusion and human exposure to trichloroethylene. J Environ Sci Health A Tox Hazard Subst Environ Eng 50, 1360–1368. 10.1080/10934529.2015.1064275.

11. Forand, S.P., Lewis-Michl, E.L., and Gomez, M.I. (2012). Adverse birth outcomes and maternal exposure to trichloroethylene and tetrachloroethylene through soil vapor intrusion in New York State. Environ Health Perspect 120, 616–621. 10.1289/ehp.1103884.

12. Shirazi, E., Hawk, G.S., Holton, C.W., Stromberg, A.J., and Pennell, K.G. (2020). Comparison of modeled and measured indoor air trichloroethene (TCE) concentrations at a vapor intrusion site: influence of wind, temperature, and building characteristics. Environmental Science: Processes & Impacts 22, 802–811. 10.1039/C9EM00567F.

13. Xie, S., and Suuberg, E. (2021). The effects of temperature and relative humidity on trichloroethylene sorption capacities of building materials under conditions relevant to vapor intrusion. J Hazard Mater 401, 123807. 10.1016/j.jhazmat.2020.123807.

14. Paciência, I., Madureira, J., Rufo, J., Moreira, A., and Fernandes Ede, O. (2016). A systematic review of evidence and implications of spatial and seasonal variations of volatile organic compounds (VOC) in indoor human environments. J Toxicol Environ Health B Crit Rev 19, 47–64. 10.1080/10937404.2015.1134371.

15. Knox, K.E., Dodson, R.E., Rudel, R.A., Polsky, C., and Schwarzman, M.R. (2023). Identifying Toxic Consumer Products: A Novel Data Set Reveals Air Emissions of Potent Carcinogens, Reproductive Toxicants, and Developmental Toxicants. Environmental Science & Technology 57, 7454–7465. 10.1021/acs.est.2c07247.

16. Health, N.I.f.O.S.a. (2018). NIOSH Pocket Guide to Chemical Hazards. In C.f.D.C.a. Prevention, ed.

17. Administration, O.S.a.H. (2021). Trichloroethylene. In D.o. Labor, ed.

18. Dorsey, E.R., Zafar, M., Lettenberger, S.E., Pawlik, M.E., Kinel, D., Frissen, M., Schneider, R.B., Kieburtz, K., Tanner, C.M., De Miranda, B.R., et al. (2023). Trichloroethylene: An Invisible Cause of Parkinson’s Disease? J Parkinsons Dis 13, 203–218. 10.3233/jpd-225047.

19. Bove, F.J., Ruckart, P.Z., Maslia, M., and Larson, T.C. (2014). Mortality study of civilian employees exposed to contaminated drinking water at USMC Base Camp Lejeune: a retrospective cohort study. Environ Health 13, 68. 10.1186/1476-069x-13-68.

20. Guehl, D., Bezard, E., Dovero, S., Boraud, T., Bioulac, B., and Gross, C. (1999). Trichloroethylene and parkinsonism: a human and experimental observation. European Journal of Neurology 6, 609–611. https://doi.org/10.1046/j.1468-1331.1999.650609.x.

21. Gash, D.M., Rutland, K., Hudson, N.L., Sullivan, P.G., Bing, G., Cass, W.A., Pandya, J.D., Liu, M., Choi, D.-Y., Hunter, R.L., et al. (2008). Trichloroethylene: Parkinsonism and complex 1 mitochondrial neurotoxicity. Annals of Neurology 63, 184–192. 10.1002/ana.21288.

22. Kochen, W., Kohlmüller, D., De Biasi, P., and Ramsay, R. (2003). The Endogeneous Formation of Highly Chlorinated Tetrahydro-ß-Carbolines as a Possible Causative Mechanism in Idiopathic Parkinson’s Disease. In Developments in Tryptophan and Serotonin Metabolism, G. Allegri, C.V.L. Costa, E. Ragazzi, H. Steinhart, and L. Varesio, eds. (Springer US), pp. 253–263. 10.1007/978-1-4615-0135-0_29.

23. Goldman, S.M., Weaver, F.M., Stroupe, K.T., Cao, L., Gonzalez, B., Colletta, K., Brown, E.G., and Tanner, C.M. (2023). Risk of Parkinson Disease Among Service Members at Marine Corps Base Camp Lejeune. JAMA Neurology. 10.1001/jamaneurol.2023.1168.

24. De Miranda, B.R., Castro, S.L., Rocha, E.M., Bodle, C.R., Johnson, K.E., and Greenamyre, J.T. (2021). The industrial solvent trichloroethylene induces LRRK2 kinase activity and dopaminergic neurodegeneration in a rat model of Parkinson’s disease. Neurobiology of disease 153, 105312–105312. 10.1016/j.nbd.2021.105312.

25. Keane, P.C., Hanson, P.S., Patterson, L., Blain, P.G., Hepplewhite, P., Khundakar, A.A., Judge, S.J., Kahle, P.J., LeBeau, F.E.N., and Morris, C.M. (2019). Trichloroethylene and its metabolite TaClo lead to degeneration of substantia nigra dopaminergic neurones: Effects in wild type and human A30P mutant α-synuclein mice. Neuroscience Letters 711, 134437–134437. https://doi.org/10.1016/j.neulet.2019.134437.

26. Liu, M., Choi, D.-Y., Hunter, R.L., Pandya, J.D., Cass, W.A., Sullivan, P.G., Kim, H.-C., Gash, D.M., and Bing, G. (2010). Trichloroethylene induces dopaminergic neurodegeneration in Fisher 344 rats. Journal of Neurochemistry 112, 773–783. https://doi.org/10.1111/j.1471-4159.2009.06497.x.

27. Sauerbeck, A., Hunter, R., Bing, G., and Sullivan, P.G. (2012). Traumatic brain injury and trichloroethylene exposure interact and produce functional, histological, and mitochondrial deficits. Experimental Neurology 234, 85–94. 10.1016/j.expneurol.2011.12.012.

28. Nolwen, L.R., Daniel, W.W., and Patrik, B. (2018). The olfactory bulb as the entry site for prion-like propagation in neurodegenerative diseases. Neurobiology of Disease 109.

29. Lucchini, R.G., Dorman, D.C., Elder, A., and Veronesi, B. (2012). Neurological impacts from inhalation of pollutants and the nose–brain connection. NeuroToxicology 33, 838–841. https://doi.org/10.1016/j.neuro.2011.12.001.

30. Cichocki, J.A., Guyton, K.Z., Guha, N., Chiu, W.A., Rusyn, I., and Lash, L.H. (2016). Target Organ Metabolism, Toxicity, and Mechanisms of Trichloroethylene and Perchloroethylene: Key Similarities, Differences, and Data Gaps. Journal of Pharmacology and Experimental Therapeutics 359, 110–110. 10.1124/jpet.116.232629.

31. Riederer, P., Foley, P., Bringmann, G., Feineis, D., Brückner, R., and Gerlach, M. (2002). Biochemical and pharmacological characterization of 1-trichloromethyl-1,2,3,4-tetrahydro-β-carboline: a biologically relevant neurotoxin? European Journal of Pharmacology 442, 1–16. https://doi.org/10.1016/S0014-2999(02)01308-0.

32. Cao, S., Theodore, S., and Standaert, D.G. (2010). Fcγ receptors are required for NF-κB signaling, microglial activation and dopaminergic neurodegeneration in an AAV-synuclein mouse model of Parkinson’s disease. Mol Neurodegener 5, 42. 10.1186/1750-1326-5-42.

33. Nair, A.B., and Jacob, S. (2016). A simple practice guide for dose conversion between animals and human. J Basic Clin Pharm 7, 27–31. 10.4103/0976-0105.177703.

34. Lewek, M.D., Poole, R., Johnson, J., Halawa, O., and Huang, X. (2010). Arm swing magnitude and asymmetry during gait in the early stages of Parkinson’s disease. Gait & Posture 31, 256–260. https://doi.org/10.1016/j.gaitpost.2009.10.013.

35. Dorsey, E.R., Greenamyre, J.T., and Willis, A.W. (2023). The Water, the Air, the Marines-Camp Lejeune, Trichloroethylene, and Parkinson Disease. JAMA Neurol. 10.1001/jamaneurol.2023.1174.

36. Anroop, B.N., and Shery, J. (2016). A simple practice guide for dose conversion between animals and human. Journal of Basic and Clinical Pharmacy 7.

37. Liu, M., Shin, E.-J., Dang, D.-K., Jin, C.-H., Lee, P.H., Jeong, J.H., Park, S.-J., Kim, Y.-S., Xing, B., Xin, T., et al. (2018). Trichloroethylene and Parkinson’s Disease: Risk Assessment. Molecular Neurobiology 55, 6201–6214. 10.1007/s12035-017-0830-x.

38. De Miranda, B.R., Castro, S.L., Rocha, E.M., Bodle, C.R., Johnson, K.E., and Greenamyre, J.T. (2021). The industrial solvent trichloroethylene induces LRRK2 kinase activity and dopaminergic neurodegeneration in a rat model of Parkinson’s disease. Neurobiol Dis 153, 105312. 10.1016/j.nbd.2021.105312.

39. Maslia, M.L., Aral, M.M., Ruckart, P.Z., and Bove, F.J. (2016). Reconstructing Historical VOC Concentrations in Drinking Water for Epidemiological Studies at a U.S. Military Base: Summary of Results. Water (Basel) 8, 449. 10.3390/w8100449.

40. Lash, L.H., Fisher, J.W., Lipscomb, J.C., and Parker, J.C. (2000). Metabolism of trichloroethylene. Environ Health Perspect 108 Suppl 2, 177–200. 10.1289/ehp.00108s2177.

41. Bradford, B.U., Lock, E.F., Kosyk, O., Kim, S., Uehara, T., Harbourt, D., DeSimone, M., Threadgill, D.W., Tryndyak, V., Pogribny, I.P., et al. (2011). Interstrain differences in the liver effects of trichloroethylene in a multistrain panel of inbred mice. Toxicol Sci 120, 206–217. 10.1093/toxsci/kfq362.

42. Chiu, W.A., Campbell, J.L., Jr., Clewell, H.J., 3rd, Zhou, Y.H., Wright, F.A., Guyton, K.Z., and Rusyn, I. (2014). Physiologically based pharmacokinetic (PBPK) modeling of interstrain variability in trichloroethylene metabolism in the mouse. Environ Health Perspect 122, 456–463. 10.1289/ehp.1307623.

43. Chiu, W.A., Okino, M.S., and Evans, M.V. (2009). Characterizing uncertainty and population variability in the toxicokinetics of trichloroethylene and metabolites in mice, rats, and humans using an updated database, physiologically based pharmacokinetic (PBPK) model, and Bayesian approach. Toxicology and Applied Pharmacology 241, 36–60. https://doi.org/10.1016/j.taap.2009.07.032.

44. Valdiviezo, A., Brown, G.E., Michell, A.R., Trinconi, C.M., Bodke, V.V., Khetani, S.R., Luo, Y.S., Chiu, W.A., and Rusyn, I. (2022). Reanalysis of Trichloroethylene and Tetrachloroethylene Metabolism to Glutathione Conjugates Using Human, Rat, and Mouse Liver in Vitro Models to Improve Precision in Risk Characterization. Environ Health Perspect 130, 117009. 10.1289/ehp12006.

45. Altmann, L., Welge, P., Mensing, T., Lilienthal, H., Voss, B., and Wilhelm, M. (2002). Chronic exposure to trichloroethylene affects neuronal plasticity in rat hippocampal slices. Environmental Toxicology and Pharmacology 12, 157–167. https://doi.org/10.1016/S1382-6689(02)00032-7.

46. Henschler, D., Romen, W., Elsässer, H.M., Reichert, D., Eder, E., and Radwan, Z. (1980). Carcinogenicity study of trichloroethylene by longterm inhalation in three animal species. Archives of Toxicology 43, 237–248. 10.1007/BF00366179.

47. Kulig, B.M. (1987). The effects of chronic trichloroethylene exposure on neurobehavioral functioning in the rat. Neurotoxicology and Teratology 9, 171–178. https://doi.org/10.1016/0892-0362(87)90095-X.

48. Mensing, T., Welge, P., Voss, B., Fels, L.M., Fricke, H.-H., Brüning, T., and Wilhelm, M. (2002). Renal toxicity after chronic inhalation exposure of rats to trichloroethylene. Toxicology Letters 128, 243–247. https://doi.org/10.1016/S0378-4274(01)00545-8.

49. De Miranda, B.R., Van Houten, B., and Sanders, L.H. (2016). Toxin-Mediated Complex I Inhibition and Parkinson’s Disease. In Mitochondrial Mechanisms of Degeneration and Repair in Parkinson’s Disease, L.M. Buhlman, ed. (Springer International Publishing), pp. 115–137. 10.1007/978-3-319-42139-1_6.

50. Mortuza, T., Muralidhara, S., White, C.A., Cummings, B.S., Hines, C., and Bruckner, J.V. (2018). Effect of dose and exposure protocol on the toxicokinetics and first-pass elimination of trichloroethylene and 1,1,1-trichloroethane. Toxicol Appl Pharmacol 360, 185–192. 10.1016/j.taap.2018.09.043.

51. Hillary, L.C., Katherine, N.W., Lucas, A.S., and Daniel, W.W. (2020). Neurochemical Organization of the Ventral Striatum’s Olfactory Tubercle. Journal of Neurochemistry 152.

52. Berg, D., Borghammer, P., Fereshtehnejad, S.M., Heinzel, S., Horsager, J., Schaeffer, E., and Postuma, R.B. (2021). Prodromal Parkinson disease subtypes - key to understanding heterogeneity. Nat Rev Neurol 17, 349–361. 10.1038/s41582-021-00486-9.

53. Chen, H., and Ritz, B. (2018). The Search for Environmental Causes of Parkinson’s Disease: Moving Forward. J Parkinsons Dis 8, S9–S17. 10.3233/JPD-181493.

54. Heintz-Buschart, A., Pandey, U., Wicke, T., Sixel-Döring, F., Janzen, A., Sittig-Wiegand, E., Trenkwalder, C., Oertel, W.H., Mollenhauer, B., and Wilmes, P. (2018). The nasal and gut microbiome in Parkinson’s disease and idiopathic rapid eye movement sleep behavior disorder. Movement Disorders 33, 88–98. https://doi.org/10.1002/mds.27105.

55. Mahlknecht, P., Seppi, K., and Poewe, W. (2015). The Concept of Prodromal Parkinson’s Disease. J Parkinsons Dis 5, 681–697. 10.3233/JPD-150685.

56. Chen, H., Wang, K., Scheperjans, F., and Killinger, B. (2022). Environmental triggers of Parkinson’s disease - Implications of the Braak and dual-hit hypotheses. Neurobiol Dis 163, 105601. 10.1016/j.nbd.2021.105601.

57. Ilieva, N.M., Wallen, Z.D., and De Miranda, B.R. (2022). Oral ingestion of the environmental toxicant trichlorethylene in rats induces alterations in the gut microbiome: relevance to idiopathic Parkinson’s disease. bioRxiv, 2022.2002.2019.481161. 10.1101/2022.02.19.481161.

58. Johnson, M.E., Stecher, B., Labrie, V., Brundin, L., and Brundin, P. (2019). Triggers, Facilitators, and Aggravators: Redefining Parkinson’s Disease Pathogenesis. Trends in Neurosciences 42, 4–13. https://doi.org/10.1016/j.tins.2018.09.007.

59. Ellenbroek, B., and Youn, J. (2016). Rodent models in neuroscience research: is it a rat race? Dis Model Mech 9, 1079–1087. 10.1242/dmm.026120.

60. Dawson, T.M., Ko, H.S., and Dawson, V.L. (2010). Genetic animal models of Parkinson’s disease. Neuron 66, 646–661. 10.1016/j.neuron.2010.04.034.

61. Gezer, A.O., Kochmanski, J., VanOeveren, S.E., Cole-Strauss, A., Kemp, C.J., Patterson, J.R., Miller, K.M., Kuhn, N.C., Herman, D.E., McIntire, A., et al. (2020). Developmental exposure to the organochlorine pesticide dieldrin causes male-specific exacerbation of alpha-synuclein-preformed fibril-induced toxicity and motor deficits. Neurobiol Dis 141, 104947. 10.1016/j.nbd.2020.104947.

62. Kochmanski, J., Kuhn, N., and Bernstein, A. (2021). Parkinson’s Disease-Associated, Sex-specific Changes in DNA Methylation at PARK7 (DJ-1), ATXN1, SLC17A6, NR4A2, and PTPRN2 in Cortical Neurons. bioRxiv.

63. Adamson, A., Buck, S.A., Freyberg, Z., and De Miranda, B.R. (2022). Sex Differences in Dopaminergic Vulnerability to Environmental Toxicants - Implications for Parkinson’s Disease. Curr Environ Health Rep 9, 563–573. 10.1007/s40572-022-00380-6.

